# Human Osteochondral Granular Extracellular Matrix (gECM) Hydrogels Drive Tissue-Specific Composition and Mechanics

**DOI:** 10.64898/2026.06.04.730235

**Authors:** Juliet O. Heye, Stephanie E. Schneider, Katie Gallagher, Shannon A. Blanco, Jeanne E. Barthold, Maxwell C. McCabe, Sean P. Maroney, Kirk C. Hansen, Michael Floren, Corey P. Neu

## Abstract

Osteochondral defects remain a major clinical challenge due to the limited regenerative capacity of cartilage and the complexity of the osteochondral interface. Here, we present a human-derived granular extracellular matrix (gECM) hydrogel platform designed for translational osteochondral repair. Using otherwise discarded human donor tissues, we developed cartilage and bone gECM hydrogels under current good manufacturing practice workflows. These materials are shear-thinning, immediately hold their form, and crosslink under physiological conditions to form stable constructs. Proteomic analysis confirmed that cartilage and bone gECM retain distinct tissue-specific biochemical signatures, while mechanical characterization demonstrated tissue-relevant stiffness, with bone gECM hydrogels exhibiting greater stiffness than cartilage gECM hydrogel. Particle packing density primarily governed viscosity, whereas tissue type contributed strongly to bulk stiffness. Together, these findings establish a scalable, human-derived gECM platform that integrates tissue-specific structural and mechanical cues, and advances a clinically translatable strategy for osteochondral repair.

## INTRODUCTION

Osteoarthritis (OA) is a prevalent and debilitating joint disease affecting over 30 million individuals in the United States alone [1]. Although cartilage degradation is a defining feature, OA is increasingly recognized as a disease of the entire joint, where biomechanical and biochemical crosstalk between cartilage and subchondral bone drives disease progression and repair outcomes [2]. Osteochondral defects remain particularly challenging to treat due to the avascularity and low cellularity of cartilage, resulting in limited intrinsic repair capacity [3]. Consequently, effective and integrative osteochondral repair remains a major unmet clinical need. Current clinical strategies are limited in both accessibility and their ability to restore functional tissue. Microfracture procedures recruit marrow-derived progenitor cells but typically yield mechanically inferior fibrocartilage [3,4]. Osteochondral allografts offer a “like-for-like” approach and are considered a clinical gold standard for focal defects. However, their use is constrained by donor availability, short shelf life, and challenges in matching tissue geometry, leading to significant tissue discard [5,6]. Cell-based therapies, including matrix-induced autologous chondrocyte implantation (MACI), require multi-stage procedures, incur high costs, and are limited by donor age and cell de-differentiation during expansion [7–9]. Collectively, these approaches fail to provide a scalable and biologically instructive solution for restoring osteochondral defects.

Extracellular matrix (ECM)-derived biomaterials have emerged as a promising strategy to address these limitations by preserving native biochemical and structural cues that regulate cell behavior [10–14]. However, intact decellularized grafts remain constrained by sourcing and geometric mismatch [5], while solubilized ECM materials often lack structural integrity and mechanical function [15,16]. Processing tissue into ECM particles provides a compelling alternative, retaining key compositional and microstructural features while enabling conformal delivery to irregular defects [13,14,17]. These particles can be incorporated into hydrogel systems to support cell signaling, tissue-specific differentiation, and enhanced mechanical performance [18–24][14,18–26].

Building on this concept, we developed granular ECM (gECM) biomaterials composed of decellularized tissue particles densely packed within a thiolated hyaluronic acid (tHA) hydrogel [12–14,17]. This approach integrates top-down and bottom-up design principles, preserving native ECM architecture while enabling injectability and tunable mechanics. At high particle packing densities, gECM exhibits percolation-driven mechanics, in which the particle network dominates bulk behavior [14]. These materials are shear-thinning and extrudable, allowing minimally invasive delivery, and subsequently crosslink under physiological conditions to form stable constructs without synthetic crosslinkers [13]. Prior work with porcine-derived gECM hydrogels demonstrated chondrocyte integration and phenotypic maintenance *in vitro* [14,17], chondrogenic responses of mesenchymal stromal cells (MSCs) to cartilage-derived matrices [12], and improved cartilage repair in preclinical *in vivo* and *ex vivo* models [13,14,27].

Despite these advances, a key barrier to clinical translation remains the integration of biological performance with scalable manufacturing. Clinical adoption requires compatibility with current good manufacturing practice (cGMP), reproducibility across donors, and alignment with clinical workflows. Here, we develop a human-derived gECM hydrogel platform using discarded cartilage and bone tissues, enabling scalable sourcing while extending the utility of donated materials. The resulting acellular, flowable material is designed for *in situ* delivery in a single surgery and positioned for a simpler regulatory pathway compared to cellular therapeutics. Processing workflows are adapted for cleanroom environments, emphasizing sterility, documentation, and quality control, while avoiding cytotoxic reagents [28] and enabling donor traceability.

In this work, we develop human-derived osteochondral gECM biomaterials using discarded donor tissues and establish cGMP-compatible processing workflows to support clinical translation. We characterize material properties and donor variability to define reproducible formulations. Ultimately, this work advances a translational human-derived biomaterial platform for osteochondral repair.

## RESULTS

### cGMP tissue processing successfully reduces DNA content and controls particle size

We previously developed a decellularization protocol for porcine tissues [17] using an acid-base viral inactivation treatment [29] that is completed within 8 hours, conducive to a contract manufacturing schedule (**Figure 1A**). We adapted this decellularization protocol and post-decellularization particularization to human cartilage and bone following Current Good Manufacturing Practices (cGMP) guidelines. We set and documented controls with personnel training, material sourcing, facilities and equipment (ISO-classified clean room), and process validation. Post-processing quality checks for this project focused on reduction of DNA content and particle size control.

**Figure 1.**
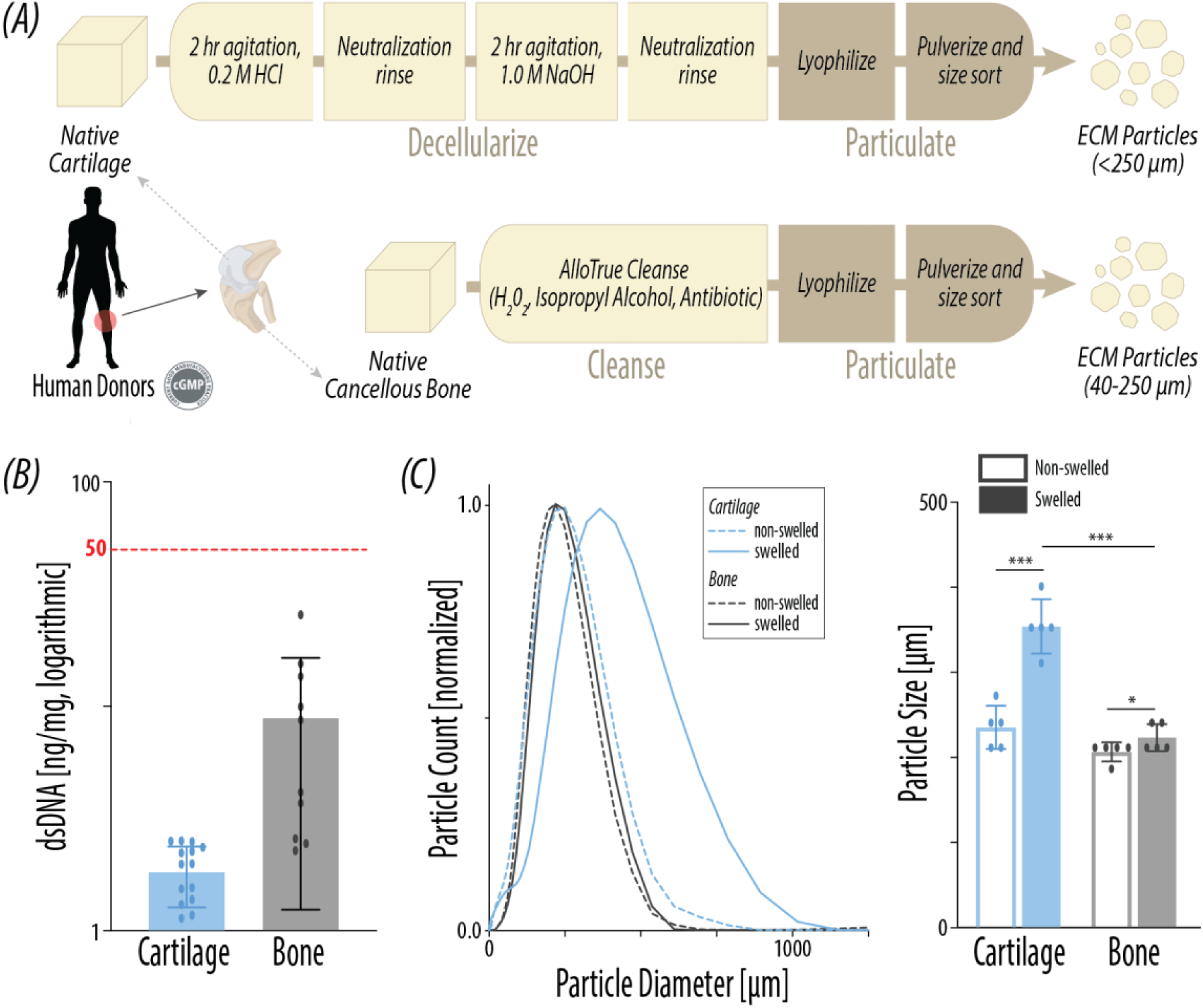
cGMP human tissue processing yields decellularized and size-sorted ECM particles. (A) Tissue is decellularized in under 8 hours in a clean room, and pulverized into <250µm (cartilage) or 40-250µm (bone) ECM particles. (B) Double-stranded DNA (N = 10-12) content after decellularization. (C) Size distribution of gECM particles in non-swelled and swelled states (N = 5). Peak particle size of gECM particles in non-swelled and swelled states (N = 5). For all plots, error bars = standard deviation, *p<0.05, **p<0.01, ***p<0.001.

Quantification of DNA content (**Figure 1B**) confirmed that residual DNA content in both human cartilage and bone was below 50 ng dsDNA per mg dry tissue [30]. Because particles are size sorted in a lyophilized form, we measured the size distribution of particles in a non-swelled state to confirm manufacturing control. Non-swelled peak sizes for both cartilage and bone fall under 250 µm. This is in line with the 250 µm sieve used to sort lyophilized particles and suggests tight control during processing. Importantly, we also characterized particle size distribution of particles in a swelled state, anticipating incorporation of particles within thiolated hyaluronic acid (tHA) and use in aqueous environments (**Figure 1C**). Swelled cartilage exhibited a significant 1.5-fold increase in size compared to its non-swelled form, while swelled cartilage only increased by 1.08-fold. These results are similar to previously characterized swelling profiles of porcine cartilage and bone [17], and are likely driven by higher proteoglycan content in cartilage [8,9,31] compared to lower proteoglycan and higher mineral content in bone [32,33]. Particle size has potential impacts on practical attributes (e.g. resolution for extrusion) and biomimetic performance (e.g. mechanics and interactions with cells).

### Cartilage and bone ECM particles retain distinct matrisome profiles

Proteomics analysis (**Figure 2A**) further confirms a reduction to <8% cellular protein content in cartilage and <1% cellular protein content in bone after processing. Proteomic analysis of matrisome components revealed that bone is primarily made up of collagens (>99%), while cartilage has ∼32% non-collagen proteins, including ∼20% proteoglycans. Heat maps showed that cartilage and bone have distinct overall proteomic profiles.

**Figure 2.**
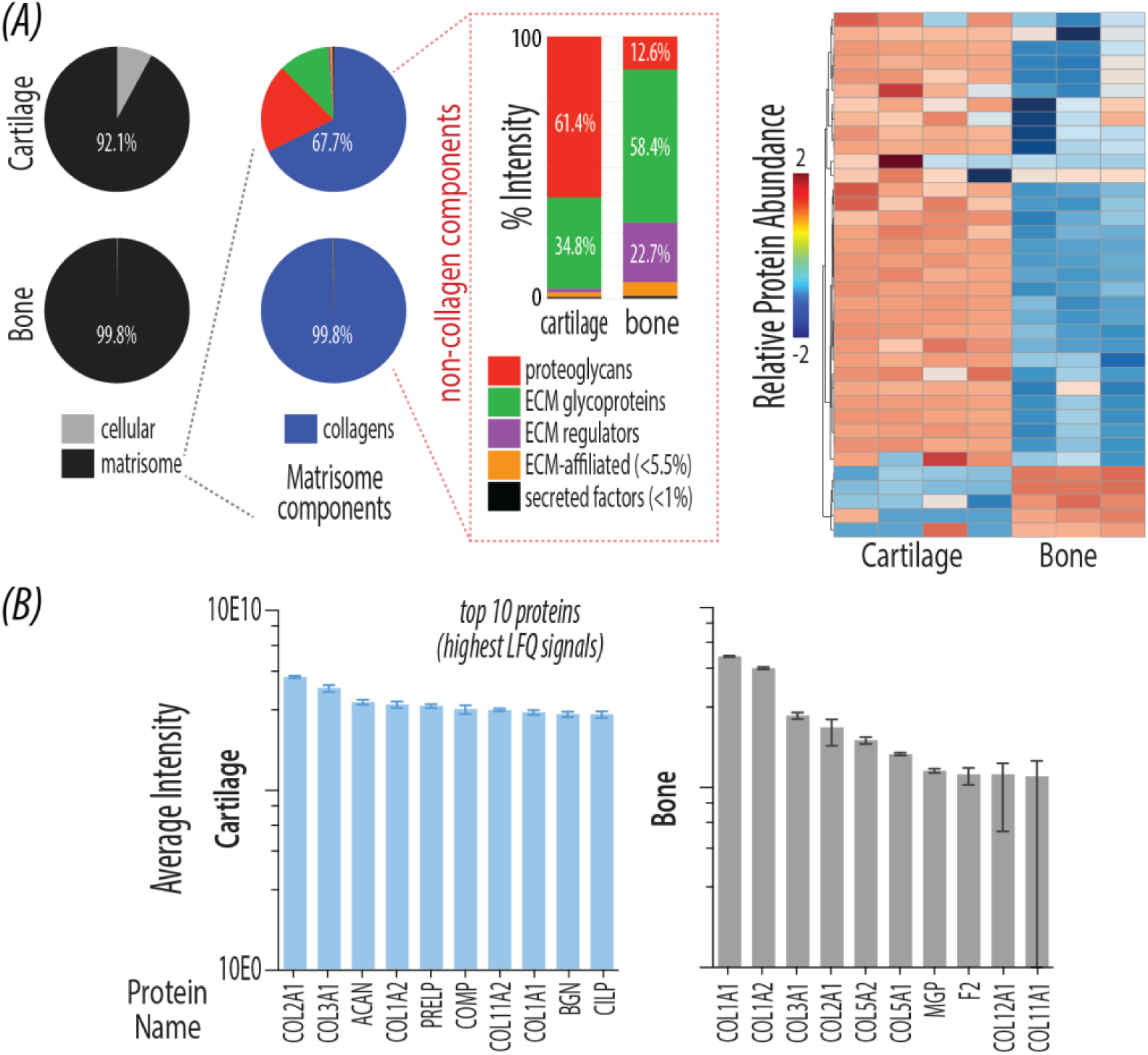
Processed human cartilage and bone ECM particles contain distinct proteomic profiles. (A) For decellularized cartilage (N = 4) and bone (N = 3), we present cellular vs. matrisomal protein content by LFQ signal. Remaining plots portray matrisome proteins only – matrisome subcategory distribution by LFQ signal, heat map with relative protein abundance, and (B) top 10 protein abundance. Error bars = standard deviation.

Notably, bone had higher abundance of collagen Type I and proteins related to collagen fibrillogenesis and mineralization (MGP, DPT, CLEC3B) compared to cartilage. These results aligned with the top 10 proteins for each tissue type (**Figure 2B**), with cartilage exhibiting COL2A1 and key cartilage proteins (ACAN, PRELP, etc) while bone exhibited various collagens (COL1A1, COL1A2, etc).

In comparison to native cartilage (**Supplementary Figure 1**), decellularized cartilage showed a reduction of non-collagen matrisome proteins. Despite this shift, decellularized cartilage retained comparable proportional distributions among the remaining non-collagen matrisome protein categories, indicating preservation of tissue-specific ECM composition. We also saw comparable abundance of the top 10 native proteins in decellularized cartilage, however the top 10 proteins based on native protein abundance (COL2A1, PRELP, COL3A1, ACAN, etc)( **Supplementary Figure 1**) differed from that of decellularized cartilage (COL2A1, COL3A1, ACAN, COL1A2, etc) (**Figure 2B**). Most notably, there are more collagens in the top 10 proteins of decellularized cartilage compared to native cartilage.

### Cartilage gECM hydrogels swell more than bone gECM hydrogels after formation

For subsequent analyses, particles and thiolated hyaluronic acid (tHA) are combined to make granular ECM hydrogels and polymerized in cylindrical molds to form constructs for testing (**Figure 3A**). Cartilage and bone gECM hydrogels maintain pH of 7-7.5 after mixing and appear white in color. The cartilage particle-to-tHA formulation was based on previous work with porcine tissues [17]. We characterized two formulations for bone: a low particle-to-tHA ratio based on previous work with porcine tissues [17] and a high ratio that achieved minimal phase separation between particles and tHA (**Supplementary Figure 2**). Cartilage gECM hydrogels polymerized in 45 mins at 37C, similar to previous methods with porcine tissues [17]. In contrast, both bone ratios required overnight polymerization (∼16hrs) to produce physically intact constructs.

**Figure 3.**
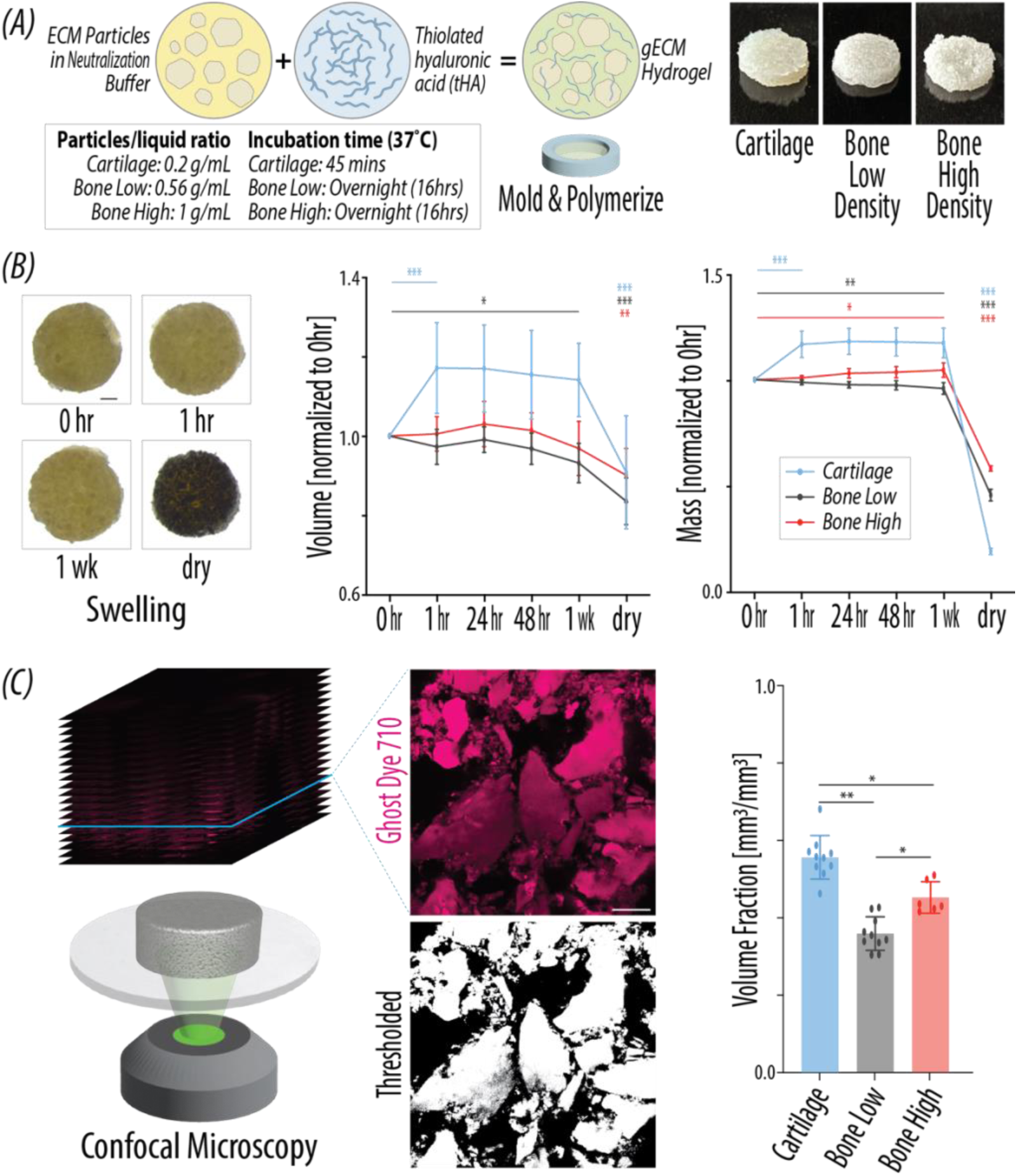
Cartilage gECM hydrogels swell more and have denser particle packing compared to bone gECM hydrogels after formation. (A) One formulation of cartilage and two formulations of bone gECM hydrogels are made by mixing thiolated hyaluronic acid (tHA) and ECM particles in neutralization buffer together. The mixed gECM hydrogel is molded in a PDMS ring and polymerized to form cylindrical constructs for characterization. Shown are representative images of polymerized samples. (B) Representative top-down images of gECM hydrogel samples during swelling, used to quantify surface area for volume calculations. Scale bar = 1mm. Mass and volume of gECM hydrogels over the course of 1 week swelling in DPBS, normalized to initial volume immediately after mixing and polymerization (N = 6-10). (C) Representative confocal slices through optical depth of polymerized and swelled cartilage gECM hydrogel sample. Selected slices were thresholded in MATLAB, and area fraction of particles was averaged across slices to calculate volume fraction of particles to total volume (N = 6-10). Scale bar = 100 µm. For all plots, error bars = standard deviation, *p<0.05, **p<0.01, ***p<0.001.

Because application of these materials involves aqueous environments (*in vitro* and *in vivo*), we measured the volume and mass of polymerized gECM constructs during swelling. Volume and mass were normalized to the constructs immediately after polymerization (0hr timepoint), to quantify changes relative to their initial form (**Figure 3B**). Cartilage plateaued in volume after 1hr, with a significant 1.17-fold increase in volume between 0hrs to 1hr. Both bone ratios maintained initial volume over 1 week, with only the low bone ratio showing a significant 0.93-fold decrease in volume between 0hr and 1wk. Both cartilage and the low bone ratio showed significant decreases in volume between 1wk and the dry form, while the high bone ratio was not significantly different between these time points. This may be due to a high content of dense mineralized bone particles resisting compaction during lyophilization. When looking at swelling mass, cartilage again increased significantly by 1.17-fold between 0hr and 1hr, indicating a mass plateau after 1hr of swelling. Both bone ratios maintained their initial mass over 1 week, with low bone showing a significant 0.96-fold decrease and high bone showing a significant 1.05-fold increase in mass between 0hr and 1wk. All gECM hydrogels showed a significant decrease in mass between 1wk and dry form, indicating water content in the hydrogels. We additionally normalized swelling parameters to dry values, report raw volume and mass, and show top-down images for all gECM formulations (**Supplementary Figure 3**). Overall, cartilage plateaus in volume and mass by 1hr, while bone effectively stays constant over 1 week. Cartilage has greater increases between initial form (0hr), fully swelled form, and dry form, while we only see modest changes in the dry form of bone. This indicates that cartilage gECM has a higher capacity for water uptake, which reflects the swelling behavior of particles alone (**Figure 1C**) and supports that particle swelling is the primary driver of overall gECM hydrogel swelling.

### gECM hydrogels are shear-thinning, and particle packing density drives viscosity

We then quantified particle-to-total volume fraction for each gECM hydrogel in their fully swelled form (**Figure 3C**). Cartilage had a significantly higher particle volume fraction compared to both bone ratios, and high bone had a significantly higher volume fraction compared to low bone, as expected. The volume fraction for cartilage matches a previously identified percolation threshold for porcine cartilage gECM [14], where particles have formed a network that drives gECM mechanics. While we have not measured a percolation threshold for bone, previous findings regarding percolation [14] suggest that the high bone ratio has a more interconnected particle network compared to the low bone ratio.

Viscosity of gECM hydrogels immediately after mixing showed shear-thinning behavior, with viscosity decreasing as shear rate increased (**Figure 4A**). We saw a significant effect of gECM type (cartilage, low bone, high bone, tHA-only) on viscosity (p<0.001). Viscosity at 10 s^−1^ was selected for comparison as a moderate shear condition relevant to material handling and injection, and before potential effects of material slipping during rheology. All gECM demonstrated significantly higher viscosity compared to tHA-only, confirming that the inclusion of ECM particles influences viscosity. Low bone was significantly lower than both cartilage and high bone, but cartilage and high bone had no statistical differences. While we saw no statistical differences in viscosity slope (0-10 s^−1^) between gECM types, we observe that HA has the lowest slope compared to gECM hydrogels. The low viscosity and shallow slope of tHA-only supports the qualitative nature of HA being already runny and minimally affected by shear. In contrast, the high viscosity and steeper slope of the gECM indicates ability to hold shape paired with injectibility.

**Figure 4.**
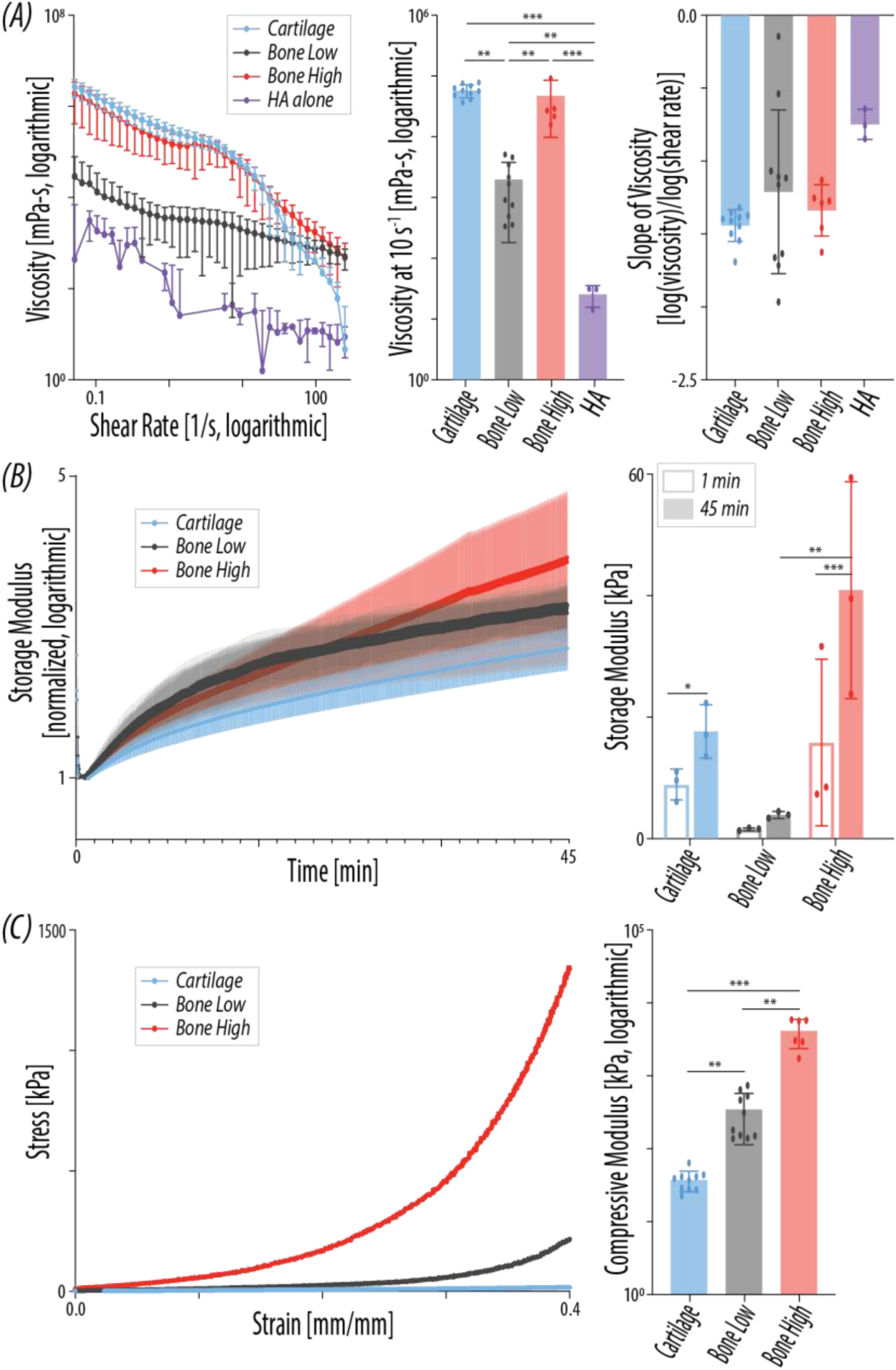
Human gECM hydrogels are shear-thinning but solid-like upon mixing, and stabilize over time, with mechanics driven by tissue type and particle packing density. (A) gECM hydrogels demonstrate shear-thinning viscosity (gECM N = 6-10, HA N = 3). Mean viscosity at 10s^−1^ shear rate. Slope of log(viscosity)/log(shear rate) between 0-10s^−1^ shear rate. Error bars = standard deviation. (B) Storage modulus of gECM hydrogels over 45 mins at 37°C, normalized to storage modulus at 1 min (N = 3). Error bars = SEM. Storage modulus (kPa) of gECM hydrogels at 1 min and 45 min at 37°C (N = 3). Error bars = standard deviation. (C) Representative stress-strain curves of gECM hydrogels during bulk unconfined compression. Bulk compressive moduli of gECM hydrogels calculated between 20-30% strain (N = 6-10). Error bars = standard deviation. For all plots, *p<0.05, **p<0.01, ***p<0.001.

Viscosity results also support a direct relationship between particle packing density and viscosity. High viscosity measurements correlate with high particle volume fraction, with cartilage exhibiting the highest volume fraction and viscosity, high bone exhibiting slightly lower volume fraction and viscosity, and low bone exhibiting the lowest volume fraction and viscosity. This was further supported by testing a medium packing density of bone gECM that exhibited viscosity between the high and low bone ratio gECM (**Supplementary Figure 2**).

### gECM hydrogels are solid-like upon mixing and continue to stabilize over time, with storage modulus influenced by particle volume fraction and tissue type

We assessed rheological properties of the gECM hydrogels immediately after mixing components for 45 minutes at 37°C (**Figure 4B**). We performed preliminary amplitude and frequency sweeps to determine oscillatory strain and frequency [34] (**Supplementary Figure 4A**), which were kept constant during the time sweep test. Loss modulus remained steady and below the storage modulus for all gECM hydrogels across the entire 45-minute test (**Supplementary Figure 4B**), suggesting that the network of ECM particles are interconnected enough to make the gECM behave like a solid even when uncured [35,36]. Qualitatively, we observe that the gECM hydrogels hold their shape immediately after mixing, which further verifies the rheological results. Storage modulus, normalized to the value at 1 minute [37] (**Figure 4B**), showed continuous stabilization over time, likely driven by thiol-crosslinking [13]. Storage modulus significantly increased between 1 and 45 mins for each gECM. Interestingly, low bone had significantly lower absolute storage modulus compared to cartilage and high bone at 1 min and 45 min. Importantly, the 45 min time point does not reflect storage modulus of physically stable bone gECM constructs, as we know they need to be at 37°C overnight to fully stabilize. In amplitude and frequency sweeps of polymerized gECM samples (cartilage after 45 min and bone after 16hr 37°C), storage modulus of low bone is greater than that of cartilage (**Supplementary Figure 4A**). Cartilage and high bone storage moduli were not significantly different at 1 min, but at 45 min, high bone was significantly higher than cartilage. This suggests that both particle volume fraction and tissue type influence storage modulus in the early time points of polymerization. Particle volume fraction primarily drives storage modulus, as seen with low bone demonstrating the lowest storage modulus in the 45 min time sweep. However, when volume fraction is comparable (e.g. cartilage and high bone), tissue type drives storage modulus (high bone > cartilage), aligning with known relative stiffnesses of these tissues. Tissue-specific particle size and shape could also modulate storage modulus under oscillatory strain [38].

### Particle volume fraction and tissue type drive bulk gECM stiffness

After polymerization and swelling, we measured bulk gECM stiffness. Representative stress-strain curves revealed a J-shape for all tissues, with lowest stress in cartilage gECM and highest stress in high bone gECM (**Figure 4C**). We highlighted the compressive moduli calculated at 20-30% strain where the effect of percolation is exaggerated, being consistent with previous work with porcine-derived gECM hydrogels [17]. This was also motivated by some bone samples reaching maximum load cell force in the 30-40% strain range. Compressive modulus calculated between 20-30% strain was significantly different between all gECM types, with cartilage exhibiting the lowest stiffness followed by low bone and then high bone. We additionally calculated compressive modulus at 4 regions of strain (0-10%, 10-20%, 20-30%, and 30-40%) (**Supplementary Figure 5**). Overall, the compressive moduli increased with increasing strain, but the relative ranking of compressive moduli (cartilage < low bone < high bone) was preserved across all regions of strain. We observed a larger difference between the stiffness of cartilage and low bone at 20-40% compared to 0-20%, likely due to compaction of particles in the low bone samples at higher strain. While we have not identified a percolation threshold for bone, previous work [14] suggests that the mechanics of the low bone ratio may have more influence from the tHA component compared to the high bone ratio. The high bone ratio has a higher volume fraction and thus likely has a more geometrically percolated particle network, leading to the particles acting as the primary driver of mechanics [14].

## DISCUSSION

In this study, we adapted our gECM technology to human tissues for application in osteochondral repair, using human donor tissues that would otherwise be discarded. We focused on manufacturing de-risking by using a decellularization process with no detergents [28], implementing cGMP-compatible workflows, and performing processing in a clean room environment. We demonstrated processing repeatability through consistent reduction of DNA content and control of particle size as key manufacturing metrics. We also demonstrated that the human bone and cartilage gECM hydrogels demonstrate important practical attributes including shear-thinning nature and ability to solidify under physiological temperature and pH. These materials exhibited differences in biomaterial composition and mechanics, including distinct proteomic profiles and bulk stiffness between cartilage and bone.

From a translational perspective, this acellular biomaterial approach offers several advantages over current surgical options. It follows a shorter path to market compared to cell-based therapies, requires only a single surgical procedure, and enables immediate filling of geometrically-varied defects with materials capable of withstanding load. The use of discarded donor tissue extends the utility of donated materials and does not rely on intact donor geometry [5]. In this study, we assessed donor variability by separating biological replicates by tissue donor, although future approaches may include pooling tissues to reduce variability. Tissue processing was performed in an ISO-classified clean room under cGMP guidelines, supporting manufacturing de-risking toward a regulatory pathway. DNA content was reduced below the industry threshold of 50 ng/mg [30], minimizing the risk of adverse immune responses. Particle size in the non-swelled state was controlled and aligned with the manufacturing process. However, the swelled particle size, which is more representative of its state when in clinical or *in vitro* use, was greater than in its non-swelled form, particularly for cartilage. Because particle size can influence extrusion resolution, volume fraction, mechanics, and cell interactions [11,14,20,22,23,49], quantifying swelling behavior enables future optimization, such as sorting highly swelling tissues to smaller diameters to achieve target sizes in the swelled state.

We developed gECM hydrogels for human cartilage and bone that fulfill practical delivery and application attributes. They are more shear-thinning and exhibit higher viscosity compared to tHA-only hydrogels, supporting their ability to be extruded while retaining shape fidelity. The relationship between storage and loss modulus further confirmed that the materials behave as solid-like immediately after mixing. This is a useful handling quality, because it suggests gECM hydrogels maintain their shape enough to be molded to variable defect shapes/sizes. Swelling behavior of the hydrogels mirrored particle swelling, suggesting that particle properties drive bulk swelling. These findings have practical implications, as bone defects may be filled to size due to minimal swelling, while cartilage defects may require slight underfilling to account for expansion. The hydrogels polymerize under physiological temperature and pH, supporting *in situ* solidification. Interestingly, bone gECM required longer polymerization times than cartilage. This may be due to fewer thiol groups on bone particles available for disulfide bonding with tHA [13], suggesting that the HA may be polymerizing around the particles more than directly to them. This is also supported by the bone gECM polymerization time aligning with tHA-only polymerization time [14].

Cartilage and bone gECM biomaterials also exhibited tissue-specific properties. The tissue particles preserved distinct proteomic signatures, highlighting their ability to deliver tissue-specific biochemical cues. Despite an overall reduction in non-collagen components following decellularization in cartilage, the maintained proportional distribution of non-collagen proteins indicates retention of key tissue-defining ECM features. The materials additionally exhibited tissue-relevant stiffnesses, with both bone formulations remaining stiffer than cartilage, suggesting that bone-derived particles are intrinsically stiffer than cartilage-derived particles and contribute directly to bulk mechanical behavior. This indicates that tissue-specific mechanical cues are preserved at both the particle and bulk levels, with potential implications for replicating *in vivo* mechanics and influencing cell behavior through mechanobiological pathways.

Particle packing density and tissue type influenced material properties in distinct ways. Viscosity profiles showed that higher packing density corresponded with higher viscosity, and this relationship was consistent across low, medium, and high bone ratios. Low packing density bone behaved more similarly to tHA, with lower viscosity and a shallower viscosity slope, while higher packing densities resulted in steeper slopes and higher viscosity. Packing density also correlates with particle volume fraction, and this trend was consistent across both bone ratios and cartilage. These results suggest that packing density, rather than tissue type, is the primary driver of viscosity, consistent with previous findings in porcine gECM systems [17]. In contrast, bulk stiffness appeared to be more strongly influenced by tissue type than packing density, although both contribute. Bone gECM exhibited higher stiffness than cartilage even at lower volume fractions, suggesting that intrinsic particle stiffness plays a significant role. Within bone, lower packing density had lower stiffness, likely due to a greater contribution from the HA component towards stiffness. Higher packing density leads to greater percolation and therefore more contribution of the particles to stiffness, as opposed to HA [14]. There is likely a lower packing density threshold below which a stiffer tissue type (like bone) would still demonstrate lower stiffness than a softer tissue type (like cartilage) at high packing density. Polymerization time also played a role in mechanical behavior. At early timepoints (within 1 hour), low bone formulations exhibited lower storage modulus than cartilage, but after overnight polymerization, storage modulus increased to exceed that of cartilage. Biomaterial characterization of both low and high bone ratios should be considered in deciding which ratio to implement in future cell and clinical studies. Importantly, both low and high bone formulations demonstrated higher mechanical stiffness compared to cartilage. While the high bone ratio has more optimal viscosity and homogenous consistency, it exhibited some clogging when extruding through a syringe, suggesting that a more intentional syringe design would need to be developed for successful injection. The low bone ratio currently optimizes injectability in addition to balancing material usage, which could be advantageous for future cell and clinical studies.

In conclusion, we developed human cartilage and bone gECM biomaterials from discarded donor tissue and established a cGMP-compatible manufacturing process that supports clinical translation. These materials exhibit practical handling characteristics, including shear-thinning behavior and polymerization under physiological conditions, and retain tissue-specific biochemical and mechanical properties. Future work will evaluate cellular responses, integration of gECM hydrogels with surrounding implant tissue, and incorporation of gECM hydrogels into advanced *in vitro* models to better implement mechanical stimulation for representation of both healthy and disease conditions. Ultimately, this platform has strong potential for recapitulating cartilage and bone tissue in regenerative applications.

## MATERIALS AND METHODS

### Tissue Particle Processing

*Tissue processing:* Human articular cartilage (N=20 human donors, **Table 1**) and cancellous bone (N=20 human donors, **Table 1**) from the knee joint were sourced from AlloSource, a human donor tissue bank, and separated by donor. All tissue processing was performed in an ISO-classified clean room, following cGMP regulations. Cartilage was cut into ∼4mm^3^ pieces and decellularized using a viral inactivation method [29]: orbital agitation at RT in 0.2M HCl for 2hrs, 1.0M NaOH for 2hrs, and rinsing in 0.02M HCl and water/DPBS until neutral (pH=7.5). Bone was pulverized and cleaned following the AlloSource AlloTrue cleanse process, including rinses in hydrogen peroxide, isopropyl alcohol, and an antibiotic solution. Cartilage and bone was then frozen, lyophilized, pulverized in a stainless-steel milling jar (Tissue Lyser III, Qiagen), and size sorted to <250 µm diameter using a sieve. Bone then underwent an additional de-fat process, including an 8hr orbital agitation in ethanol (50%, 75%, 95%, 100% for 2hrs each) [51], and overnight orbital agitation in DPBS. It was then lyophilized and sorted to include particles 40-250µm diameter. Cartilage and bone tissue particles were packaged in sterile vials, k-pack, and chevron packaging until use.

**Table 1.** Age and sex (Male, Female) of human adMSC and human cartilage/bone donors, grouped for cell studies.

| adMSC Donor | Cartilage Donor | Bone Donor |
| --- | --- | --- |
| 23M | 39F, 39M | 54F, 72M |
| 22F | 38F, 38M | 20F, 16M |
| 23M | 39F, 39M | 39F, 37M |
| 28F | 38F, 36M | 49M, 68M |
| 30M | 38F, 39M | 24M, 33M |
| 23M | 39M, 38M | 32M, 51M |
| 23M | 38F, 38M | 42M, 65M |
| 21F | 36F, 39M | 57M, 66M |
| 21M | 38M, 36M | 56M, 64M |
| 23F | 38F, 36M | 61M, 76M |

### Tissue Particle Characterization

#### DNA content

Double-stranded DNA (dsDNA) was extracted from processed cartilage and bone tissue particles (Zymo DNA Extraction Kit) and quantified using a PicoGreen assay (ThermoFisher Invitrogen).

#### Particle Size and Swelling

Particle size distribution of gECM particles suspended in 100% ethanol (non-swelled) or DI water (swelled) was measured in a laser diffraction particle size analyzer (Mastersizer 3000). Mie theory was applied to calculate volume-based particle size distributions. Measurements were considered valid when residual error was <1%. The Beer-Lambert law was used to monitor optical obscuration and confirm appropriate sample concentration. Tissue-specific refractive indices were used (cartilage: 1.40 [52], bone: 1.55 [53]), while the absorptive index (0.1) and density (1.00 g/cm^3^) were held constant across tissues [54–58].

#### Proteomics sample preparation

Lyophilized native cartilage, decell cartilage, and decell bone samples were processed using a single-fraction hydroxylamine (1 M NH_2_OH−HCl, 4.5 M Gnd−HCl, 0.2 M K_2_CO_3_, pH adjusted to 9.0 with NaOH) extraction to isolate insoluble ECM (iECM) components. Extraction buffers were applied at a ratio of 200 μL/mg of the starting tissue dry weight (0.5–2 mg), and homogenized at power 8 for 3 min. Samples were subsequently incubated at 45°C with shaking (1000 rpm) for 4 h. Following incubation, the samples were spun for 15 min at 18,000 x g, and the supernatant was removed and stored as the iECM fraction. All collected fractions were immediately frozen and stored at −80°C until further proteolytic digestion. Extracted proteins were subjected to enzymatic digestion overnight (16 h) at 37°C with trypsin (1:100 enzyme to protein ratio) using a filter aided sample preparation (FASP) approach as previously described [59] and desalted during Evotip loading.

#### LC-MS/MS analysis

Digested peptides (200 ng) were loaded onto individual Evotips following the manufacturer‘s protocol and separated on an Evosep One chromatography system (Evosep, Odense, Denmark) using a Pepsep column, (150 µm inter diameter, 15 cm) packed with ReproSil C18 1.9 µm, 120Å resin. Samples were analyzed using the instrument default “30 samples per day” LC gradient. The system was coupled to the timsTOF Pro mass spectrometer (Bruker Daltonics, Bremen, Germany) via the nano-electrospray ion source (Captive Spray, Bruker Daltonics). The mass spectrometer was operated in PASEF mode. The ramp time was set to 100 ms and 10 PASEF MS/MS scans per topN acquisition cycle were acquired. MS and MS/MS spectra were recorded from m/z 100 to 1700. The ion mobility was scanned from 0.7 to 1.50 Vs/cm^2^. Precursors for data-dependent acquisition were isolated within ± 1 Th and fragmented with an ion mobility-dependent collision energy, which was linearly increased from 20 to 59 eV in positive mode. Low-abundance precursor ions with an intensity above a threshold of 500 counts but below a target value of 20000 counts were repeatedly scheduled and otherwise dynamically excluded for 0.4 min.

#### Global proteomic data analysis

Data was searched using MSFragger v4.3 via FragPipe v23.1 [60]. Precursor tolerance was set to ±15 ppm and fragment tolerance was set to ±25 ppm. Data was searched against UniProt restricted to *Homo sapiens* with added common contaminant sequences [61] (46,127 total sequences, downloaded 9/25/25). Enzyme cleavage was set to semi-specific trypsin for all samples. Fixed modifications were set as carbamidomethyl (C). Variable modifications were set as oxidation (M), oxidation (P) (hydroxyproline), deamidation (NQ), Gln->pyro-Glu (N-term Q), and acetyl (protein N-terminus). Label free quantification was performed using IonQuant v1.11.11 with match-between-runs enabled and default parameters. Results were filtered to 1% FDR at the peptide and protein level. Matrisome protein annotations were derived from MatrisomeDB [62] (annotation levels 1 and 2), in addition to in-house generated annotations for matrisome subcategories (annotation levels 3-5).

### gECM hydrogel Formulation

#### Hyaluronic Acid Functionalization

Glucoronate carboxyl groups on hyaluronic acid (Lifecore Biomedical, HA-100K-5) were replaced with thiol groups following previously established protocols to produce thiolated hyaluronic acid (tHA) [14], with purification via tangential flow filtration. The substitution rate was confirmed to be 18%–23% (thiolated mmols/unthiolated mmols) using a standard Ellman’s assay (Ellman’s solution, ThermoFisher).

#### gECM hydrogel Mixing

The gECM hydrogel formulations consisted of 10 mg/mL tHA packed with 0.2 g/mL (cartilage) [14], and 0.56 g/mL (low bone) [17] or 1 g/mL (high bone) lyophilized particles. For subsequent characterization of polymerized samples, gECM hydrogels were molded into cylindrical constructs (1.5 mm height x 5 mm diameter) and incubated for 45 min (cartilage) or overnight (∼16hours, bone) at 37°C.

### gECM hydrogel Biophysical Characterization

#### Swelling

Polymerized gECM hydrogel constructs were incubated in DPBS at room temperature for 1 week and then lyophilized. Mass and volume were measured immediately after polymerization (0 hr), at 4 timepoints during incubation in DPBS (1 hr, 24 hrs, 48 hrs, 1 wk), and after lyophilization. Volume was obtained by measuring height with calipers and calculating surface area from thresholded (Image J) stereoscopic images (Leica).

#### Particle Volume Fraction

Tissue particles in polymerized constructs were stained with Ghost Dye 710 (Cytek Biosciences) and imaged via confocal microscopy (Nikon A1R, 20x objective, NA=0.75). We obtained three-dimensional 5µm-step z-stack images at 3 different locations per sample. Using a custom MATLAB code, we selected, thresholded, and averaged the area over 15µm depth to calculate particle volume to total construct volume.

### gECM hydrogel Mechanical Characterization

#### Rheology

Within 5 minutes of mixing the gECM hydrogels, a room temperature rheological shear sweep (0.1 to 250 s^−1^) measuring viscosity was performed (MCR 702, Anton Paar). Separately, we performed a 45-minute time sweep measuring storage (G’) and loss (G’’) modulus at 37°C, with constant oscillation at 0.5% shear strain and 0.9 rad/s frequency (MCR 702, Anton Paar). Parameters were determined by identifying linear trends in preliminary amplitude (0.01-100% strain, 1 rad/s frequency) and frequency (0.01-100 rad/s, 1% strain) sweeps for gECM hydrogels in both uncured (immediately after mixing) and cured (after 45mins at 37°C) states. Rheological tests were performed with 8mm parallel plates and a 3mm (cartilage and high bone) or 2mm (low bone) gap height [34].

#### Bulk Compression

Polymerized gECM hydrogel constructs were compressed (unconfined) to 40% strain at a quasi-static 0.1%/s strain rate (MCR 702, Anton Paar). Compressive modulus was calculated between 0-10%, 10-20%, 20-30%, and 30-40% strain [14].

### Statistical Analysis

Mixed-model analyses of variance (ANOVAs) were performed on linear mixed models with tissue type and treatment (e.g. time, swell state, etc) as the co-factors, donor (tissue donor and/or cell donor) as random effects, and the resulting measurement variable as the response (DNA, particle size, swelling volume, swelling mass, volume fraction, viscosity, storage modulus, compressive modulus, surface area, volume). All residuals were checked, and if a non-normal distribution was identified, the data was transformed for statistical analysis. Post hoc multiple comparisons were performed using Tukey’s honestly significant difference (HSD) correction. Significance was defined as p<0.05.

## SUPPLEMENTARY FIGURES

**Supplemental Figure 1.**
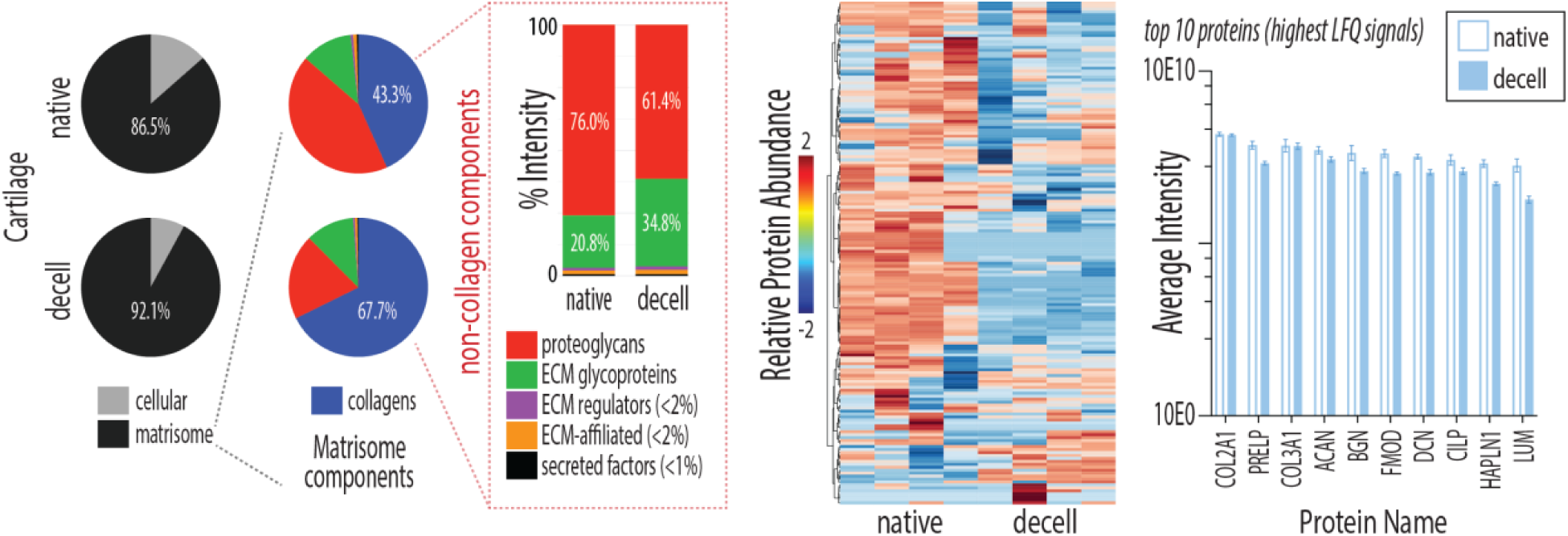
Decellularization of ECM particles retains key matrisome components, including collagens. Cartilage (N = 3) ECM particles proteomics compared before (Native) and after (Decell) decellularization. We present cellular vs. matrisomal protein content by LFQ signal. Remaining plots portray matrisomal proteins only – matrisome subcategory distribution by LFQ signal, heat map with relative protein abundance, and top 10 protein abundance. Error bars = standard deviation.

**Supplemental Figure 2.**
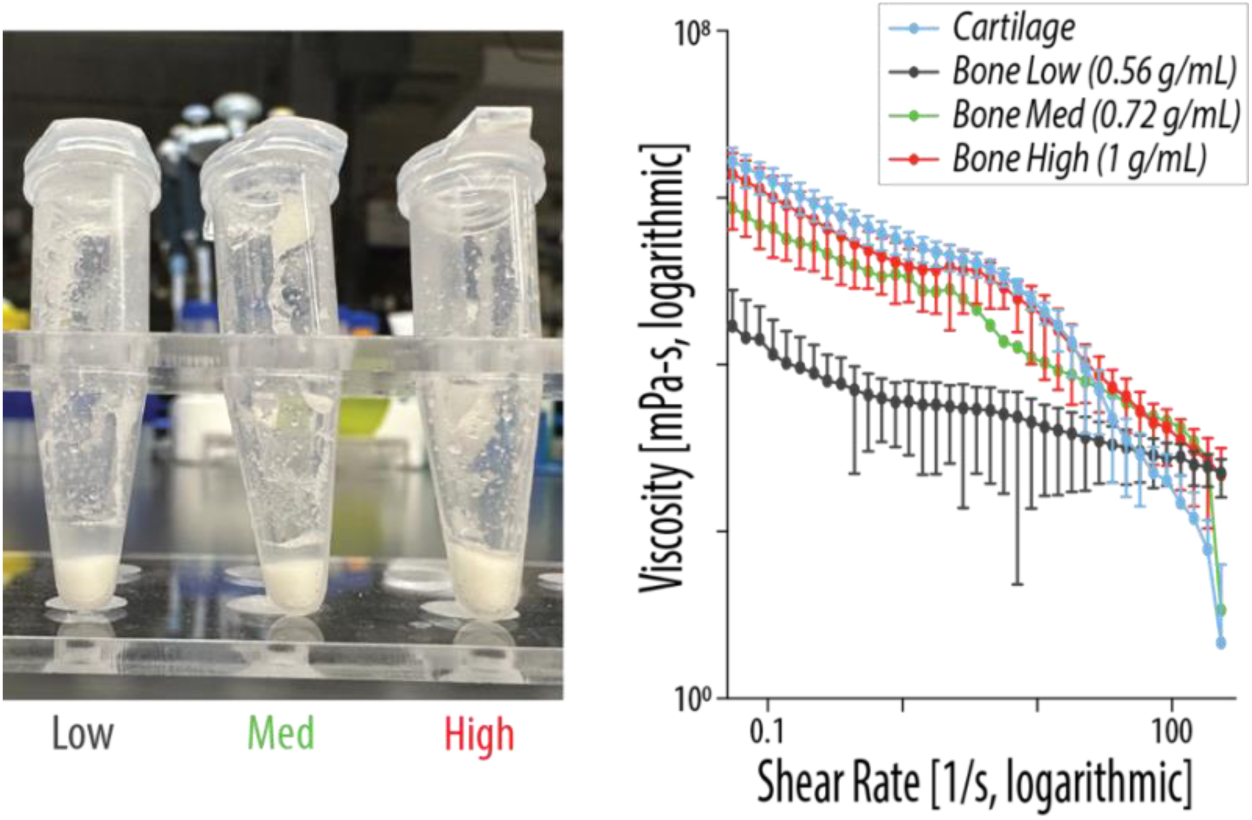
Bone particle packing density formulations. Representative images of low (0.56 g/mL), medium (0.72 g/mL), and high (1 g/mL) bone gECM hydrogel formulations before polymerization, after particle settling. Viscosity of cartilage gECM (0.2 g/mL) and 3 bone gECM formulations (Cartilage, low bone, high bone N = 6-10; med bone N = 1). Error bars = standard deviation.

**Supplemental Figure 3.**
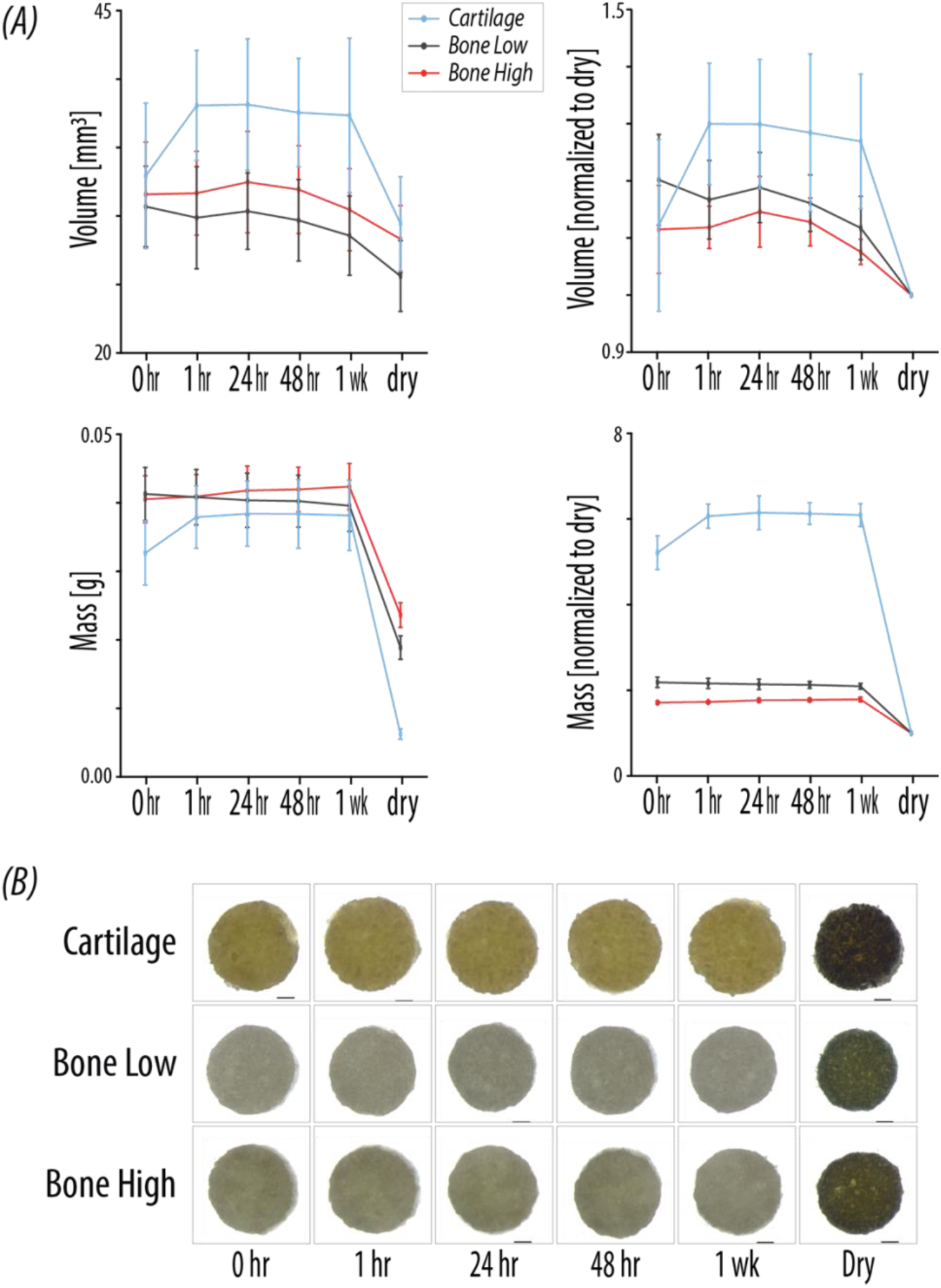
Swelling data raw values and normalized to dry. (A) Volume and mass for gECM hydrogels over 1 week swelling in PBS. Shown here as raw values, and values normalized to the dry volume or mass. Error bars = standard deviation. (B) Representative top-down images of gECM hydrogels during swelling, used to quantify surface area for volume calculations.

**Supplemental Figure 4.**
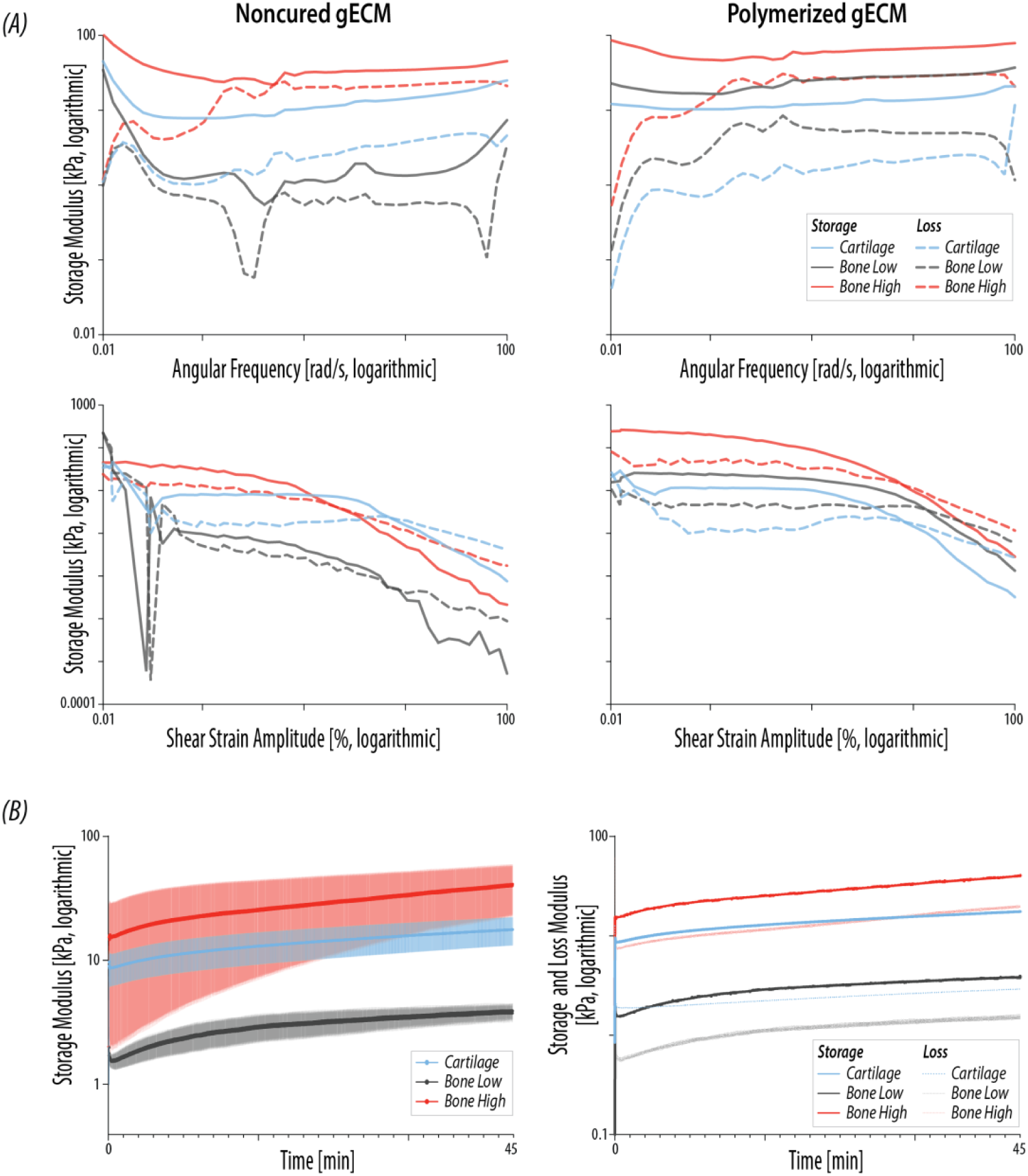
Rheological amplitude, frequency, and time sweeps. (A) Amplitude and frequency sweeps for gECM hydrogels in a noncured (immediately after mixing) and polymerized (cartilage incubated 45 mins and bone incubated ∼16hrs at 37°C) state (N = 3-5). (B) Raw storage modulus over a 45-minute time sweep (N = 3). Error bars = SEM. Raw mean storage and loss modulus over a 45-minute time sweep (N = 3).

**Supplemental Figure 5.**
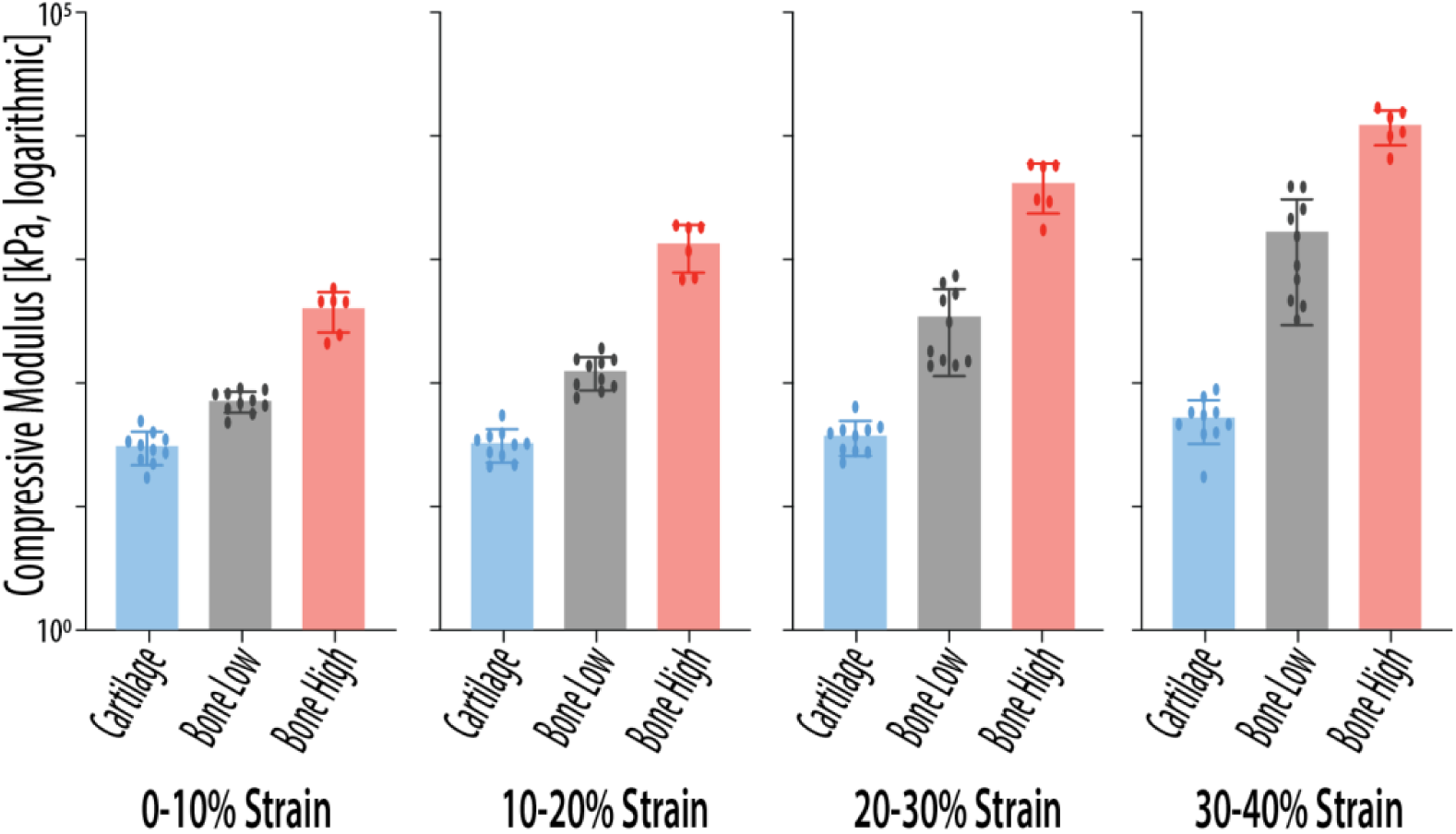
gECM hydrogel compressive modulus calculated for various strain windows. Compressive modulus calculated at 0-10%, 10-20%, 20-30%, and 30-40% strain for gECM hydrogels (N = 6-10). Error bars = standard deviation.

## Acknowledgments

The authors acknowledge AlloSource for providing human tissues and a cGMP clean room for tissue processing.

## Funding

The authors gratefully acknowledge funding from the following sources: National Institutes of Health U01 AR082845 and R01 AR083379 (CPN), National Institutes of Health P30CA06934 funded Mass Spectrometry Proteomics Shared Resource [RRID SCR_021988], and National Science Foundation Graduate Research Fellowship (JOH).

## Authors

Juliet O. Heye, Stephanie E. Schneider, Katie Gallagher, Shannon A. Blanco, Jeanne E. Barthold, Maxwell C. McCabe, Sean P. Maroney, Kirk C. Hansen, Michael Floren, Corey P. Neu

## Author contributions

Conceptualization: JOH, SES, JEB, MF, CPN

Methodology: JOH, SES, KG, SAB, JEB, MCM, SPM, KCH, CPN

Visualization: CPN, JOH

Funding acquisition: CPN, JOH, SES, JEB

Supervision: CPN, SES

Writing – original draft: JOH

Writing – review & editing: All authors

## Competing interests

Authors JOH, JEB, and CPN have equity in TissueForm, Inc. JEB and CPN are co-inventors on a filed patent pertaining to the material used in the manuscript: (US 18/039,242, Particulate materials for tissue mimics).

## Notes

### Summary of Updates

Changed the author contributions (added co-first authorship)

